# Leveraging FracMinHash Containment for Genomic *d*_*N*_/*d*_*S*_

**DOI:** 10.1101/2025.11.12.688019

**Authors:** Judith S. Rodriguez, Mahmudur Rahman Hera, David Koslicki

## Abstract

Increasing availability of genomic data demands algorithmic approaches that can efficiently and accurately conduct downstream genomic analyses. These analyses, such as evaluating selection pressures within and across genomes, can reveal developmental and environmental pressures. One such commonly used metric to measure evolutionary pressures is based on the ratio of non-synonymous and synonomous substitution rates, *d*_*N*_/*d*_*S*_. Conventionally, the *d*_*N*_/*d*_*S*_ ratio is used to infer selection pressures employing alignments to estimate total non-synonymous and synonymous substitution rates along protein-coding genes. However, this process can be time consuming and not scalable for larger datasets. Recently, a fast, approximate similarity measure, FracMinHash containment, was introduced and related to average nucleotide identity. In this work, we show how FracMinHash containment can be used to quickly estimate *d*_*N*_/*d*_*S*_ enabling alignment-free estimations at a genomic level.

Through simulated and real world experiments, our results indicate that employing FracMinHash containment to estimate *d*_*N*_/*d*_*S*_ is scalable, enabling pairwise *d*_*N*_/*d*_*S*_ estimations for 85,205 genomes within 5 hours. Furthermore, our approach is comparable to traditional *d*_*N*_/*d*_*S*_ methods, representing sequences subject to positive and negative selection across various mutation rates. Moreover, we used this model to evaluate signatures of selection between Archaeal and Bacterial genomes, identifying a previously unreported metabolic island between *Methanobrevibacter sp*. RGIG2411 and *Candidatus Saccharibacteria bacterium* RGIG2249. We present, FracMinHash *d*_*N*_/*d*_*S*_, a novel alignment-free approach for estimating *d*_*N*_/*d*_*S*_ at a genome level that is accurate and scalable beyond gene-level estimations while demonstrating comparability to conventional alignment-based *d*_*N*_/*d*_*S*_ methods. Leveraging the alignment-free similarity estimation, FracMinHash containment, pairwise *d*_*N*_/*d*_*S*_ estimations are facilitated within milliseconds, making it suitable for large-scale evolutionary analyses across diverse taxa. It supports comparative genomics, evolutionary inference, and functional interpretation across both synthetic, and complex biological datasets.

**Availability and implementation:** A version of the implementation is available at https://github.com/KoslickiLab/dnds-using-fmh.git. The reproduction of figures, data, and analysis can be found at https://github.com/KoslickiLab/dnds-using-fmh_reproducibles.git.

**Supplementary information:** Supplementary data are available at PLOS Computational Biology online.

**Author summary:** Understanding how evolution shapes genomes helps us learn about the pressures organisms face in their environments. Scientists traditionally measure this by comparing genetic changes that alter proteins versus those that don’t, a ratio that reveals whether natural selection is preserving or changing genes. However, this conventional approach requires computationally intensive sequence alignments, making it impractical for analyzing the massive genomic datasets now available. We developed a faster, alignment-free method to estimate evolutionary pressure across entire genomes. Our approach uses a computational technique called FracMinHash that compresses genomic information while preserving meaningful patterns. We tested our method on both simulated and real-world data, including over 85,000 microbial genomes, completing the analysis in just five hours whereas traditional methods would take days or weeks for the same analysis. The results were comparable to traditional methods and correctly identified genes under different types of selection. Using this approach, we discovered a previously unreported shared genetic region between an archaeal and bacterial species from the goat gut microbiome, suggesting ancient gene transfer between these distant branches of life. Our method makes large-scale evolutionary analysis practical for diverse applications, from tracking microbial strains to understanding adaptation in complex microbial communities, potentially accelerating discoveries in comparative genomics and evolutionary biology.

## 1 Introduction

The strength and direction of substitution events shape the coding regions of a genome and can be inferred by measuring selection pressures [1, 2]. Selection pressure of coding regions is measured by the ratio of codon substitutions occurring at non-synonymous to synonymous sites (*d*_*N*_*/d*_*S*_), denotated as *ω* [2, 3, 4, 5, 6], and is used to understand evolutionary events that influence biophysical activities such as transcript degradation, protein stability, [7], and protein function. Non-synonymous substitutions are changes in a codon that alter the resulting amino acid in a protein sequence, whereas synonymous substitutions are changes in a codon that do not affect the resulting amino acid. In this ratio, the rate of non-synonymous to synonymous substitutions can indicate the pressure of selection that is being exerted on a protein-coding sequence [8]. The inference of selection pressures is based on the amount of substitutions that affect the protein sequence. An excessive amino acid change along a gene is considered to be under adaptive changes (*d*_*N*_*/d*_*S*_ >1), conversely, the preservation of amino acids is considered as constraining selection(*d*_*N*_ */d*_*S*_ < 1).

Traditional *d*_*N*_/*d*_*S*_ programs have been developed for gene level estimations. A variety of models employing different algorithms and codon substitution models to account for mutations that influence selection have been developed. Earlier approaches relied on approximation methods. For example, LWL85 considered transition and transversion mutations to avoid the overestimation of synonymous sites and the underestimation of non-synonymous sites [9]. Later, LBP93 modified LWL85 ignoring the ratio of transition and transversion mutations [10]. Subsequent models for *d*_*N*_/*d*_*S*_ incorporated maximum-likelihood approaches such as GY94 [11], NY98 [12], and YN00 [13], as well as bayesian methods, like Omegamap [14] and CodonRates [15]. While these methods are typically a straightforward process that involves sequence alignments, they can become computationally demanding when applied on a plethora of genes across thousands of genomes. Estimating selection can be further challenged when applied on a metagenomic sample, where instead of estimating selection pressures of a single genome, estimations are made on a mixture of genomes from diverse organisms, making the estimation of *d*_*N*_/*d*_*S*_ particularly demanding due to the intensive processing time required for alignments and model inference.

Consequently, these conventional methods of estimating *d*_*N*_/*d*_*S*_ ratios may not scale for the vast number of genomic sequences present in metagenomic datasets, limiting the utility of *d*_*N*_/*d*_*S*_ to smaller genetic scales, such as individual genes [2] becoming a bottleneck in the measurement and analysis of selection pressures. Although often limited to smaller genetic datasets, *d*_*N*_/*d*_*S*_ pipelines have been developed to study entire genomes, yielding insights into the selection pressures related to recombination [16], genome size [17, 18], antimicrobial resistance [19], carcinomas [20]. Only recently have approaches for genomic level analyses been developed.

GenomegaMap was developed to address this problem by utilizing parent-dependent mutation models to avoid the use of product of approximate conditionals [5]. An extension of the earlier *d*_*N*_/*d*_*S*_ estimator OmegaMap [14], GenomegaMap was developed to consider recombination as a phylogeny-free approach by avoiding the inference of multiple unrooted tree topologies that can be produced while reducing computational demands. Although innovative, GenomegaMap requires sequence alignments similar to earlier models.

In contrast, recently popular methods of genomic analysis include so-called *k*-mer sketching techniques, which create lower-dimensional representations of high-dimensional *k*-mer profiles, enabling alignment-free approaches scalable to tasks such as *d*_*N*_/*d*_*S*_ estimations. One such method, called FracMinHash, utilizes a fraction of *k*-mers obtained from pairs of metagenomic sequences to quickly estimate a similarity index called the containment. Recently, the containment index has been statistically tested and related to the simple mutation model [21, 22]. Building on this model, average nucleotide identity (ANI) can be estimated from FracMinHash containments, thereby enabling the estimation of nucleotide mutation rates between genome sequences [22]. Derived from the containment, approximations of ANI are facilitated to estimate differences of a pair of genomes. Adapting this framework, the same theory can be extended to estimating the mutation rates of protein sequences using the average amino acid identity (AAI). By assuming a simple mutation model, ANI and AAI can be calculated from FracMinHash containments facilitating estimations of *d*_*N*_/*d*_*S*_ and inference of selection pressures at the genomic level.

Using an alignment-free method to estimate genomic *d*_*N*_/*d*_*S*_ to analyze and interpret evolutionary pressures within a microbial community can be used to quickly provide insights into gene loss [17], genome sizes [18], species delineation [23] and evolving speeds [24], as well as drug discovery and development [25]. Therefore, we present *FracMinHash d*_*N*_/*d*_*S*_, an alignment-free approach to estimating *d*_*N*_/*d*_*S*_ at a genomic level and its application for metagenomic data.

## 2 Methods

In the following section, we describe the theoretical approach to estimating *d*_*N*_/*d*_*S*_ at a genomic level. To provide context, we first recall the following key definitions.

### 2.1 Preliminaries

Recall the definition of a FracMinHash sketch of a set *A* with scale factor 0 ≤ *s* ≤ 1 and a uniform hash function *h* with range [0, *H*] [26]:

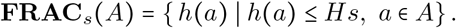

The debiased Fractional Containment Index [22] is an unbiased estimator of the true containment index:

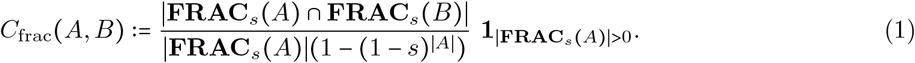

The equation for sequence similarity estimation (ANI or AAI) from FracMinHash Containment is reproduced here for completeness. See [22] for these and other analytical details.

### 2.2 FracMinHash *d*_*N*_*/d*_*S*_: ratio of non-synonymous to synonymous mutations

The calculation for *d*_*N*_/*d*_*S*_ is defined as the ratio between non-synonymous and synonymous substitution rates, denoted by *dN* and *dS*, respectively. *dN* is defined as the ratio of the total number of non-synonymous mutations *pN* and the total number of non-synonymous sites *N* ; whereas *dS* is defined as the total number of synonymous mutations *pS* and the total number of synonymous sites *S*. Using these notations, *d*_*N*_/*d*_*S*_ is defined as follows.

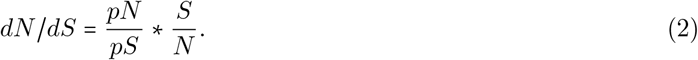

We first make our simple mutation model explicit to estimate *d*_*N*_/*d*_*S*_ using FracMinHash. Assume we are given a nucleotide sequence *W* = *W*_1_⋯*W*_*n*_ with the open reading frame starting at *W*_1_*W*_2_*W*_3_. The mutation process can be described by position-wise Bernoulli random variables: *X*_1_, *X*_2_, *X*_3_ through *X*_*n*_, where every *X*_*i*_ ∼ Bernoulli(*p*) for 1 ≤ *i* ≤ *n*. Considering the reading frame, we know the codons of *W* consist of (*W*_1_, *W*_2_, *W*_3_), (*W*_4_, *W*_5_, *W*_6_), etc. Let *Z*_*i*_ be a random variable that indicates if the *i*^th^ codon has a nucleotide mutation in it:

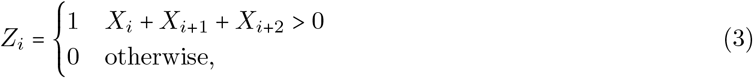

and let *N*_*i*_ be an indicator that, under the mutation model, the *i*-th codon of *W* has experienced a non-synonymous mutation:

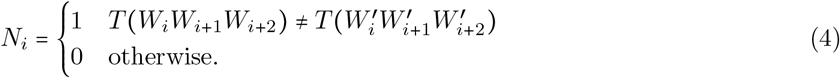

Hence, Pr[*N*_*i*_ = 1] is equal to the observed AAI. Let us call this value *p*_*AA*_ := Pr[*N*_*i*_ = 1]. Note that a non-synonymous mutation can only happen when a nucleotide mutation has occurred, so Pr[*N*_*i*_ = 1∣*Z*_*i*_ = 0], and the absence of a nucleotide mutation implies a non-synonymous mutation cannot happen: Pr[*N*_*i*_ = 0∣*Z*_*i*_ = 0] = 1. Thus, we can calculate that

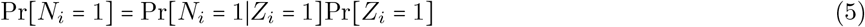

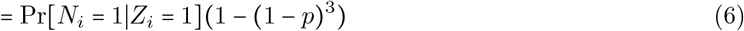

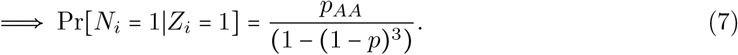

Similarly, we calculate that

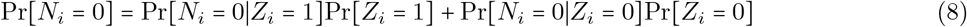

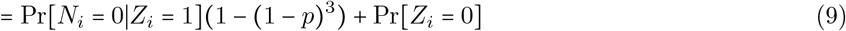

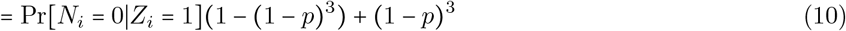

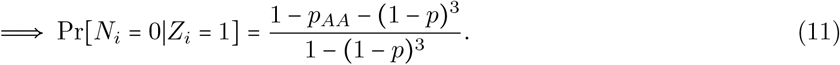

Thus, we can estimate the ratio of expected non-synonymous to expected synonymous mutations as:

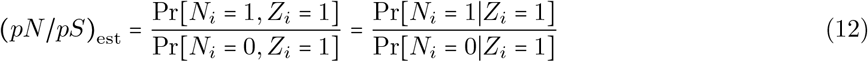

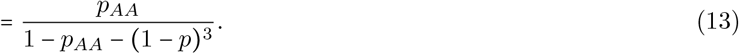

If *k*_*nt*_-mers are used to sketch the nucleotide sequences, then the nucleotide mutation rate *p* is estimated as follows [22]:

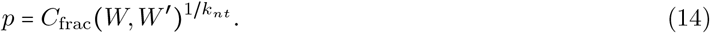

Similarly, if *k*_*aa*_-mers are used to sketch the amino-acid sequences, then the amino-acid mutation rate *p*_*AA*_ is estimated as follows [22]:

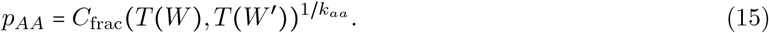

Next, we turn to estimating *S* / *N* . If we define *r* to be the total number of codons, then the total number of sites (including both the synonymous and the non-synonymous sites) is simply 3*r*. If *S* is the total number of synonymous sites, then the total number of non-synonymous sites is simply 3*r* − *S*. As suggested by the 1986 Nei and Gojobori *d*_*N*_/*d*_*S*_ model (NG86), on average, there are 0.72 synonymous sites in the first positions of the codons, and there are 0.05 synonymous sites in the third positions of the codons [27]. Therefore, *S* = 0.72*r* + 0.05*r* = 0.77*r*, and *N* = 3*r* − 0.77*r* = 2.23*r*. Using these observations, we estimate *S* / *N* as follows:

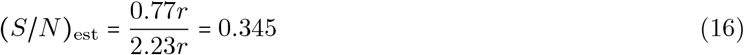

Using Equations 13, 14, 15, and16, we estimate *d*_*N*_/*d*_*S*_ as follows:

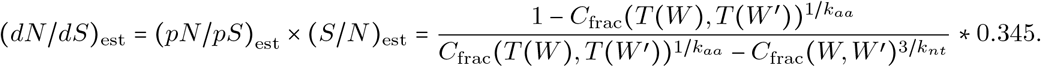

### 2.3 Implementation of FracMinHash *d*_*N*_*/d*_*S*_

To describe the theoretical implementation of *FracMinHash d*_*N*_/*d*_*S*_, we provide a workflow diagram in **Figure 1**). This is an alignment-free, efficient, and accurate approach to estimating genomic-level *d*_*N*_/*d*_*S*_ by leveraging the *k*-mer-based comparative tool sourmash [26] to generate FracMinHash sketches and calculate containment indices. Implemented in Python, *FracMinHash d*_*N*_/*d*_*S*_ depends on sourmash for efficient compression and comparison of sequence datasets of varying sizes. Specifically, it utilizes branchwater, a sourmash plugin, enabling rapid, low-memory estimation of pairwise FracMinHash-based *d*_*N*_/*d*_*S*_.

**Figure 1:**
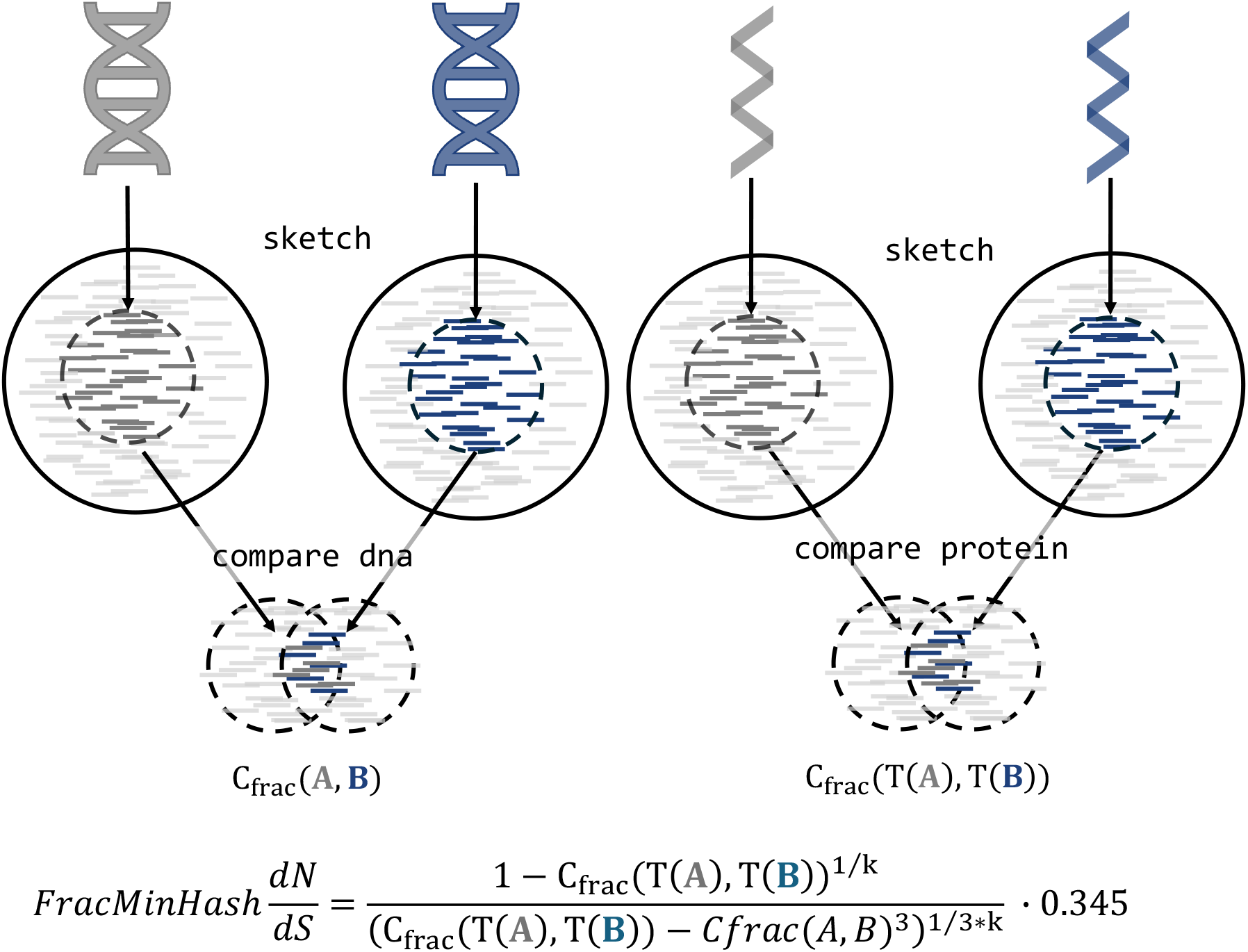
*FracMinHash d*_*N*_/*d*_*S*_ pipeline. *FracMinHash d*_*N*_/*d*_*S*_ employs sourmash to sketch genomic datasets and perform pairwise comparisons to obtain FracMinHash containments that are used in esitmating FracMinHash *d*_*N*_/*d*_*S*_. (a) Transcriptomic and proteomic information of genomes are taken as input to (b) sketch a scaled *k*-mer representation of the genomic information. (c) The skteches are compared between DNA and protein to get FracMinHash containments, respectively. (d) The estimated FracMinHash *d*_*N*_/*d*_*S*_ is calculated from FracMinHash containment of DNA and protein.

In the intitial steps of estimating FracMinHash *d*_*N*_/*d*_*S*_, sourmash is used to sketch genomic datasets and conduct pairwise comparisons, producing FracMinHash containment indices required for estimating FracMinHash *d*_*N*_/*d*_*S*_. The command *sourmash scripts manysketch* simultaneously generates signatures for DNA and protein sequences using the appropriate *k*-size parameters (i.e., *k*_*aa*_ == 3 ∗ *k*_*nt*_). After generating these sketches, the pairwise containment indices for both DNA and protein sequences are calculated individually using the command *sourmash scripts multisearch*. Finally, these containment indices are utilized within Python scripts to estimate FracMinHash-based *d*_*N*_/*d*_*S*_. All relevant code and documentation are publicly available at dnds-using-fmh GitHub repository.

## 3 Results

### 3.1 Robust and accurate *d*_*N*_*/d*_*S*_ estimations demonstrated on synthetic data

#### Random positive and negative selected simulations

A key insight interpreted by *d*_*N*_/*d*_*S*_ values is the ability to infer selection—whether positive (*d*_*N*_/*d*_*S*_ > 1) or negative (*d*_*N*_/*d*_*S*_ < 1) selection is being observed. We assess the accuracy to which selection can be interpretted by *FracMinHash d*_*N*_/*d*_*S*_ by simulating both positive and negative selection. This simulation is based on the suggestion that most synonymous substitutions occur in the first and third position of a codon (**Supplemental Figure ??A**) [27]. We adopt this suggestion to simulate a random nucleotide sequence that is 10,002 base pairs (bp) in length and select for positive and negative selection by randomly generating a reference sequence, and introducing random mutations at a mutation rate of *p* to produce mutated query sequences.

Negative selected query sequences generated from the reference sequence are created by applying the *p*-rate to the third position of each codon. Conversely, for positive selected query sequences, the same *p*-rate was applied to the first and second positions of each codon. Results are displayed using a scatter plot where negative and positive selection have *d*_*N*_/*d*_*S*_ < 1 and *d*_*N*_/*d*_*S*_ > 1, respectively (**Supplemental Figure ??B**). Hence, 100 simulated sequences were generated for each selection pressure—positive and negative selection—to benchmark against a ground truth.

*d*_*N*_/*d*_*S*_ models are based off of codon substitution models and can be categorized as approximation [9, 27, 10], maximum likelihood [11, 28, 13], and bayesian [14, 5] models. Among approximation models, we use NG86 as our ground truth, a model that uses the Jukes-Cantor Correction in which substitution rates and nucleotide frequencies are equal [27]. Since our model is based on NG86, we use it as the ground truth to estimate *d*_*N*_/*d*_*S*_ of simulated datasets and evaluate *FracMinHash d*_*N*_/*d*_*S*_.

Estimates from this ground truth were assessed on whether the correct selection can be inferred under varying *p*-rates (0.1, 0.01, and 0.001). Across these *p*-rates, a distinction can be made between positive and negative selected datasets, suggesting that our method to simulate selection was accomplished. As expected, the sequences substituted for positive selection resulted in values greater than 1 and the negative selected sequences below 1 (**Supplemental Figure ??A**). More specifically, a *p*-rate of 0.1 would result in FracMin-Hash containments approaching 0, leading to undefined *d*_*N*_/*d*_*S*_ estimations. On the other hand, a smaller *p*-rate of 0.001 may produce sequences too similar to the reference, resulting in FracMinHash containments approaching 1, biasing *d*_*N*_/*d*_*S*_ estimations and preventing to reliably distinguish between positive and negative selection. Therefore, we continue evaluating the performance of *FracMinHash d*_*N*_/*d*_*S*_ using a *p*-rate of 0.01.

Given that the most optimal inference of selection can be achieved using a *p*-rate of 0.01, we employed sequences simulated at this rate to evaluate the usage of the *k*-size parameter in *FracMinHash d*_*N*_/*d*_*S*_ for different sequence lengths to evaluate how *k*-size and length can impact FracMinHash *d*_*N*_/*d*_*S*_ estimations when compared to a traditional *d*_*N*_/*d*_*S*_ model like NG86. Across varying *p*-rates, increasing *k*-size can considerably reduce the expected containment to 0 [22]. Thus, sequences should be sufficiently long that the *k*-size does not affect containment, enabling effective use for *FracMinHash d*_*N*_/*d*_*S*_ and avoid undefined *d*_*N*_/*d*_*S*_ estimates.

Using the selection simulation method, we generated positive and negative selected sequences with a *p*-rate of 0.01 for nucleotide sequence lengths of 5,001, 10,002 and 20,001, and estimated FracMinHash *d*_*N*_/*d*_*S*_ using *k*_*aa*_ of 5, 7, 15, and 21. Comparing FracMinHash *d*_*N*_/*d*_*S*_ estimates to the ground truth, NG86, results support that longer sequences become better correlated to the ground truth and that as *k*-size increases across different lengths, the correlation weakens (**Figure 2A**). As sequence length increases, FracMinHash *d*_*N*_/*d*_*S*_ estimations become more correlated to the traditional NG86 model for both positive and negative selected sequences. Moreover, across different lengths, *k*_*aa*_ of 5 and 7 yield the highest correlation to the ground truth with a Pearson R correlation of at least 0.90. However, at a *k*_*aa*_ of at least 15, correlations across different sequence lengths begin to decline. Overall, these results suggest that *FracMinHash d*_*N*_/*d*_*S*_ is capable of (1) accurately estimating selection for both negative and positive values and (2) providing greater resolution for longer sequences, indicating that our alignment-free FracMinHash *d*_*N*_/*d*_*S*_ model can be applicable for longer sequence, potentially at the genomic level.

**Figure 2:**
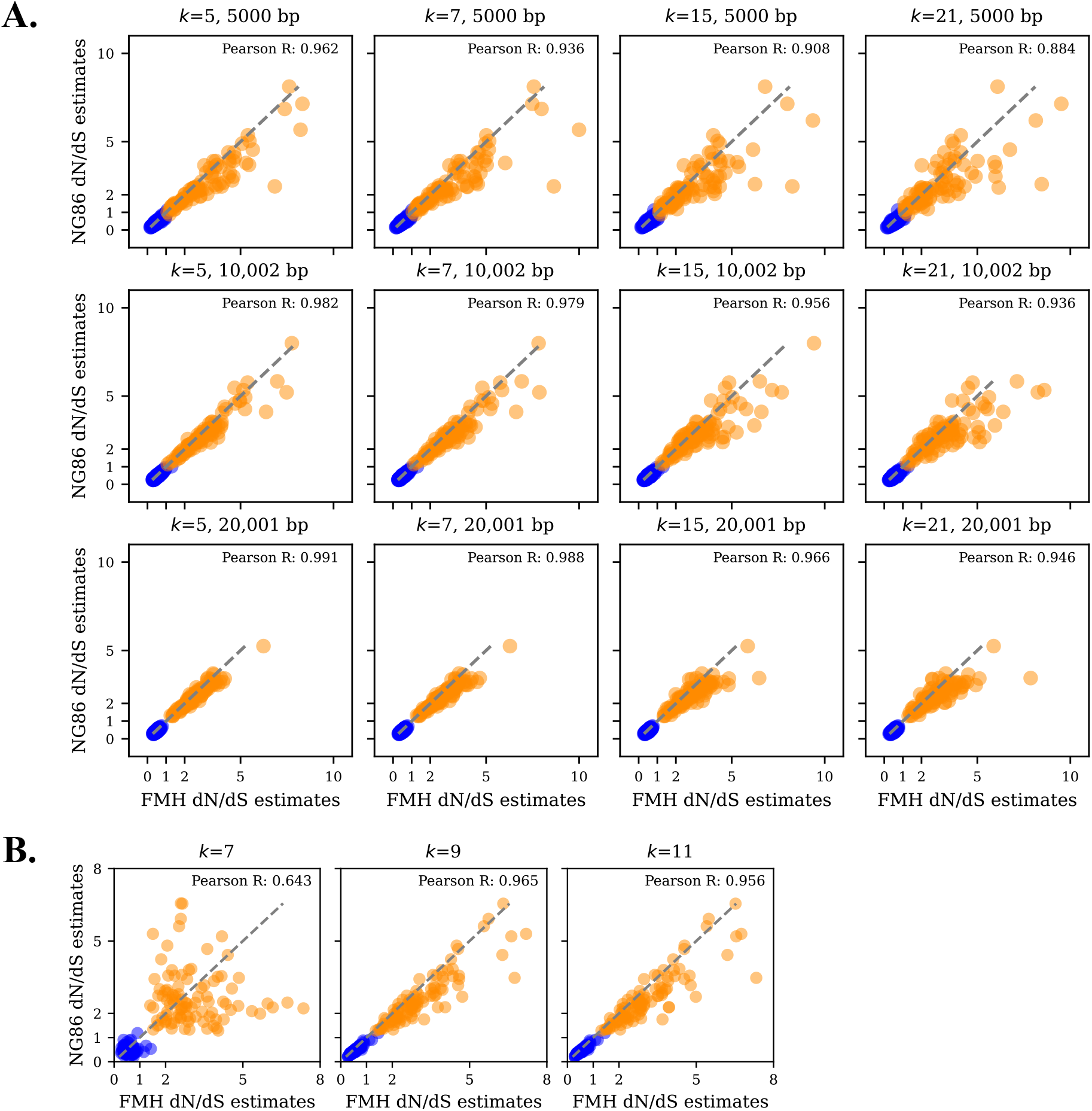
Usefulness and performance of *FracMinHash d*_*N*_/*d*_*S*_. The blue and orange data points represent the negative and positive selected simulations applied, respectively. Additionally, Pearson R correlations between dN/dS estimations by NG86 and *FracMinHash d*_*N*_/*d*_*S*_ have been included. A) Simulation applied on a random nucleotide sequence of varying lengths (i.e., 5,001bp, 10,002bp, and 20,001bp) across *k*_*aa*_-sizes 5, 7, 15, and 21. B) Simulation applied on a real gene, LAMA3, using *k*_*aa*_-sizes 7, 9, and 11.

#### Positive and negative simulations of endogenous biological sequences

Next, we want to evaluate the reliability of *FracMinHash d*_*N*_/*d*_*S*_ when a *p*-rate of 0.01 is applied on endogenous biological sequences to simulate positive and negative selection. Furthermore, we look at how varying *k*-sizes and scale factors can impact FracMinHash *d*_*N*_/*d*_*S*_ estimations. Because we observe that better resolution is achieved with longer sequences, we test *FracMinHash d*_*N*_/*d*_*S*_ on a long gene sequence and a genomic sequence. Our results have suggested that *k*_*aa*_-sizes between 5 and 15 are the most appropriate. Therefore, we utilized *k*_*aa*_-sizes 7, 9, and 11 to assess *FracMinHash d*_*N*_/*d*_*S*_ when mutations are introduced into native biological sequences. Moreover, using a scale factor of 1 (i.e., all *k*-mers are present in the sketch) indicates that *FracMinHash d*_*N*_/*d*_*S*_ is a reliable estimator to infer selection pressures; however, when applying *FracMinHash d*_*N*_/*d*_*S*_ on metagenomic datasets, a user will want to use larger scale factors to compress the data and reduce computational time. Increasing the scale factor can influence the resolution of the biological data impacting FracMinHash *d*_*N*_/*d*_*S*_ estimates; therefore, we test how larger scale factors may impact FracMinHash *d*_*N*_/*d*_*S*_ estimates of genomic sequences.

We first apply *FracMinHash d*_*N*_/*d*_*S*_ to a long gene sequence that has been selectively mutated, LAMA3, a sequence that is >10k nucleotides and generated 100 queries for both negative and positive selection. Consistent with our mutation simulation on a random sequence, we observe that positive and negative selection queries are moderately distributed across different *k*-sizes (**Figure 2B**). Positively selected sequences have a *d*_*N*_/*d*_*S*_ > 1, while most negatively selected sequences show a *d*_*N*_/*d*_*S*_ < 1, suggesting *FracMinHash d*_*N*_/*d*_*S*_ is capable of distinguishing selection pressure. Once again, we rely on the ground truth, NG86, to assess *d*_*N*_/*d*_*S*_ estimations made by *FracMinHash d*_*N*_/*d*_*S*_. A strong correlation to this traditional *d*_*N*_/*d*_*S*_ model is observed at *k*_*aa*_-sizes of 9 and 11, with a Pearson R of 0.965 and 0.956, respectively. Although, the correlation decreases when using a *k*_*aa*_-size of 7 (Pearson R=0.643), positive and negative selection remains distinguishable.

The scale factor is an important parameter that is used to compress a *k*-mer set while preserving its meaningful heterogeneity. However, there is a tradeoff when increasing the scale factor. The computational time of estimating similarity index, such as the containment, can decrease but the resolution of the *k*-mer profile can also decrease. Therefore, we assess how different scale factors impact FracMinHash *d*_*N*_/*d*_*S*_ estimations by applying our selection simulation on the genomic coding sequences of *Escherichia coli*. We use the Genome Taxonomy Database (GTDB) representative for *E. coli*, GB GCA 021307345.1, a sampled genome that is 4.4Mb with 4K annotated genes. Using a *k*_*aa*_-size of 9 at scale factors of 10, 100, and 1000, we evaluated whether selection can be distinguished as the scale factor increases. Thus, increasing the scale factor from 10 to 1000 may result in some overlap of positive and negative selection, but estimations remain distinguishable, suggesting increasing the scale factor would not have a grave impact on inference (**Supplemental Figure ??**).

We have applied *FracMinHash d*_*N*_/*d*_*S*_ on mutated endogenous biological data to assess its performance of varying *k*-sizes and scale factors. As *k*-sizes increases, we can observe robust results when compared to the ground truth NG86 with negative selected sequences being overestimated. We also tested how scale factor can impact estimations and inference, and found that even with increasing the scale factor at 1000, inferences can still be made. Our findings motivate the applicability of *FracMinHash d*_*N*_/*d*_*S*_ on real-world genomic data.

### 3.2 Efficient and reliable pairwise selection inference performed when applying *FracMinHash d*_*N*_*/d*_*S*_ on 85k genomes

#### Computational efficiency

To demonstrate the applicability of *FracMinHash d*_*N*_*/d*_*S*_ at a genomic level, we employ representative genomes of the GTDB. Specifically, we tested *FracMinHash d*_*N*_/*d*_*S*_ on coding and protein sequences of 85,205 GTDB representative genomes (Release 08-RS214; 28th April 2023), encompassing bacterial and archaeal genomes. We evaluate the performance in terms of disk usage (**Figure 3A**) and computational time (**Figure 3B**) at each step of the pipeline, which include sketching, containment estimations, and FracMinHash *d*_*N*_/*d*_*S*_ estimations, across two parameters—*k*-size and containment threshold (*t*).

**Figure 3:**
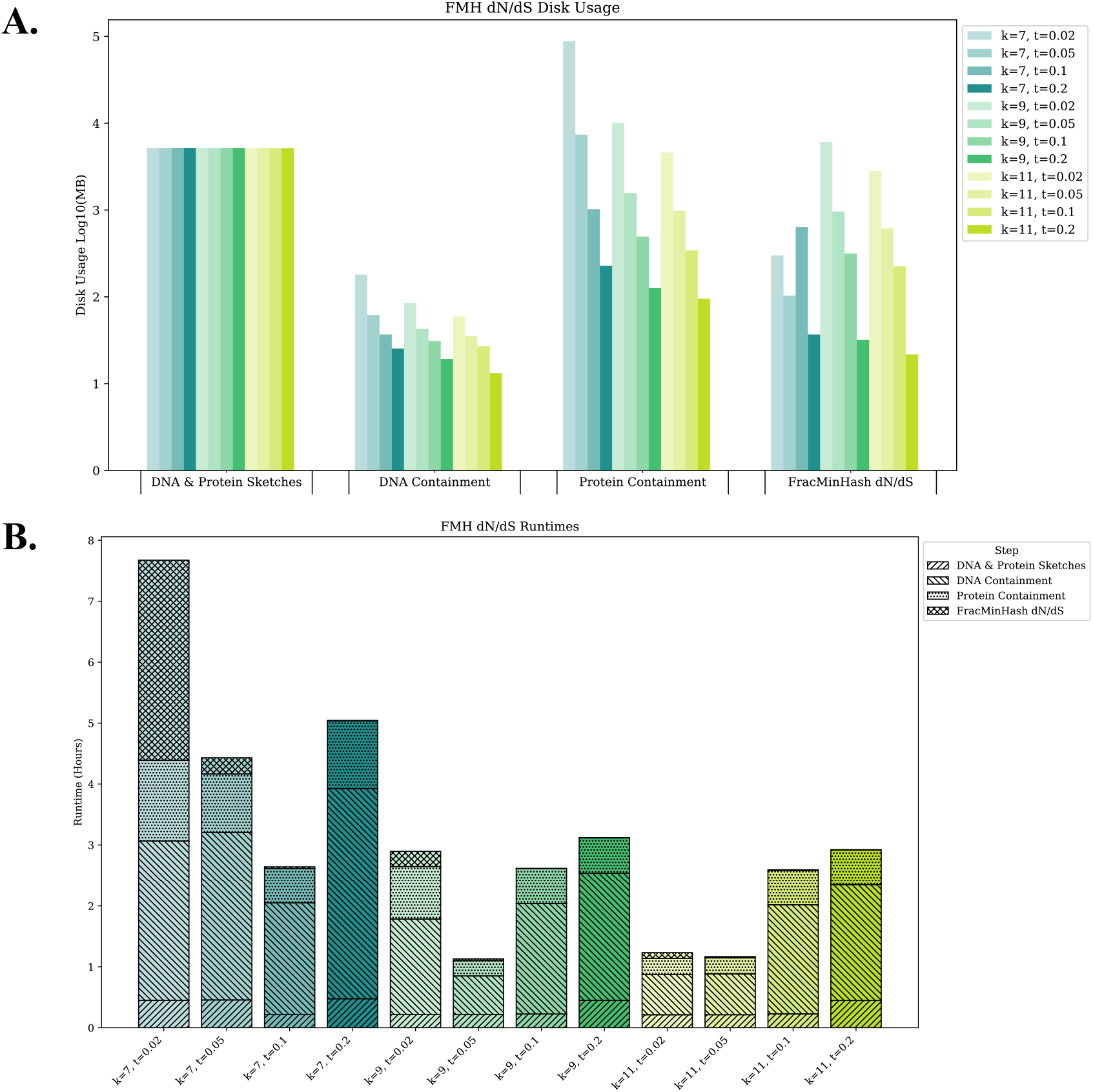
Time and disk usage evaluation across different containment thresholds. A) Disk usage description. B) Line plot represents different k-sizes (7, 9, and 11) and containment thresholds (0.02, 0.05, 0.1, and 0.2) at a scale factor of 500 can estimate pairwise FracMinHash *d*_*N*_/*d*_*S*_ for 85,205 representative genomic coding and protein sequences between 2 to 9 hours on a server with 128 CPUs

In the first step, the input FASTA files containing DNA and protein sequences are simultaneously sketched across *k*_*aa*_-sizes 7, 9, and 11, each with a scale factor of 500. Disk usage remained consistent across different *k*-sizes, and the sketching process took less than an hour to complete. The next two steps involve estimating pairwise containments for DNA and protein individually, with DNA containments performed first. We evaluate disk usage and computational time across different containment thresholds: 0.02, 0.05, 0.1, and 0.2. Since the same containment threshold is applied to both DNA and protein comparisons, greater variability is observed at the DNA level than at the protein level due to synonymous substitutions. Computationally, estimating pairwise DNA containments take longer than protein containments because DNA sketches, which are based on the codon level of an amino acid sequence (i.e., 1aa=3bp), generate more hash values than protein sketches. Additionally, as the *k*-size increases, fewer hash values are produced resulting in reduced disk usage and computational time for both DNA and protein pairwise containment estimations. The final step in the pipeline estimates the FracMinHash *d*_*N*_/*d*_*S*_. This step is generally quick, with the exception of using a *k*_*aa*_-size of 7 and containment threshold of 0.02, which required more time.

Overall, the total pipeline runtime typically ranges between 3.5 and 4 hours. Using a *k*_*aa*_-size of 7 across different containment thresholds takes the longest time. Since larger *k*-sizes generate fewer hash values from *k*-mers, increasing the *k*_*aa*_-size to 11 can dramatically decrease the time required for DNA containment estimation and overall pipeline execution. Therefore, *FracMinHash d*_*N*_/*d*_*S*_ efficiently estimates pairwise FracMinHash *d*_*N*_/*d*_*S*_ across various *k*-sizes and containment thresholds, with the total runtime varying between 1 and 8 hours. Specifically, using a *k*_*aa*_-size larger than 7 with a containment threshold of 0.05 results in systematic disk usage and computational time demonstrating that *FracMinHash d*_*N*_/*d*_*S*_ is scalable for large-scale analyses such as metagenomic data.

#### Selection estimation and inference

To contextualize these estimations to real-world evolutionary analyses, we compare our FracMinHash *d*_*N*_/*d*_*S*_ estimations to published genomic-level pairwise *d*_*N*_/*d*_*S*_ values from the GTDB database [18]. In this study, *d*_*N*_/*d*_*S*_ values were estimated using a maximum-likelihood model implemented in CodeML, a phylogenetic tool within the PAML suite [29, 18]. The median *d*_*N*_/*d*_*S*_ values within each genus were then calculated and compared to genomic sizes to evaluate associations regarding gene loss [18]. To assess whether *FracMin-Hash d*_*N*_/*d*_*S*_ can reproduce similar interpretations, we replicated the analysis by calculating the median FracMinHash *d*_*N*_/*d*_*S*_ estimations to obtain a genus-level estimates and compare results to the published dataset.

We first explore how well correlated our estimations are to those made by Martinez-Gutierrez et al., (2022) under varying *k*-sizes and containment thresholds. We find that across different *k*-sizes and containment thresholds, our estimations are being overestimated (**Figure 4A**). The most robust results occur when using a *k*_*aa*_-size of 7 showing moderate correlations across different containment thresholds. As the containment threshold increases, this correlation weakens. Therefore, a reduction in sample size will occur, as fewer genome pairs meet the threshold, thereby decreasing the statistical power when comparing *d*_*N*_/*d*_*S*_ estimations.

**Figure 4:**
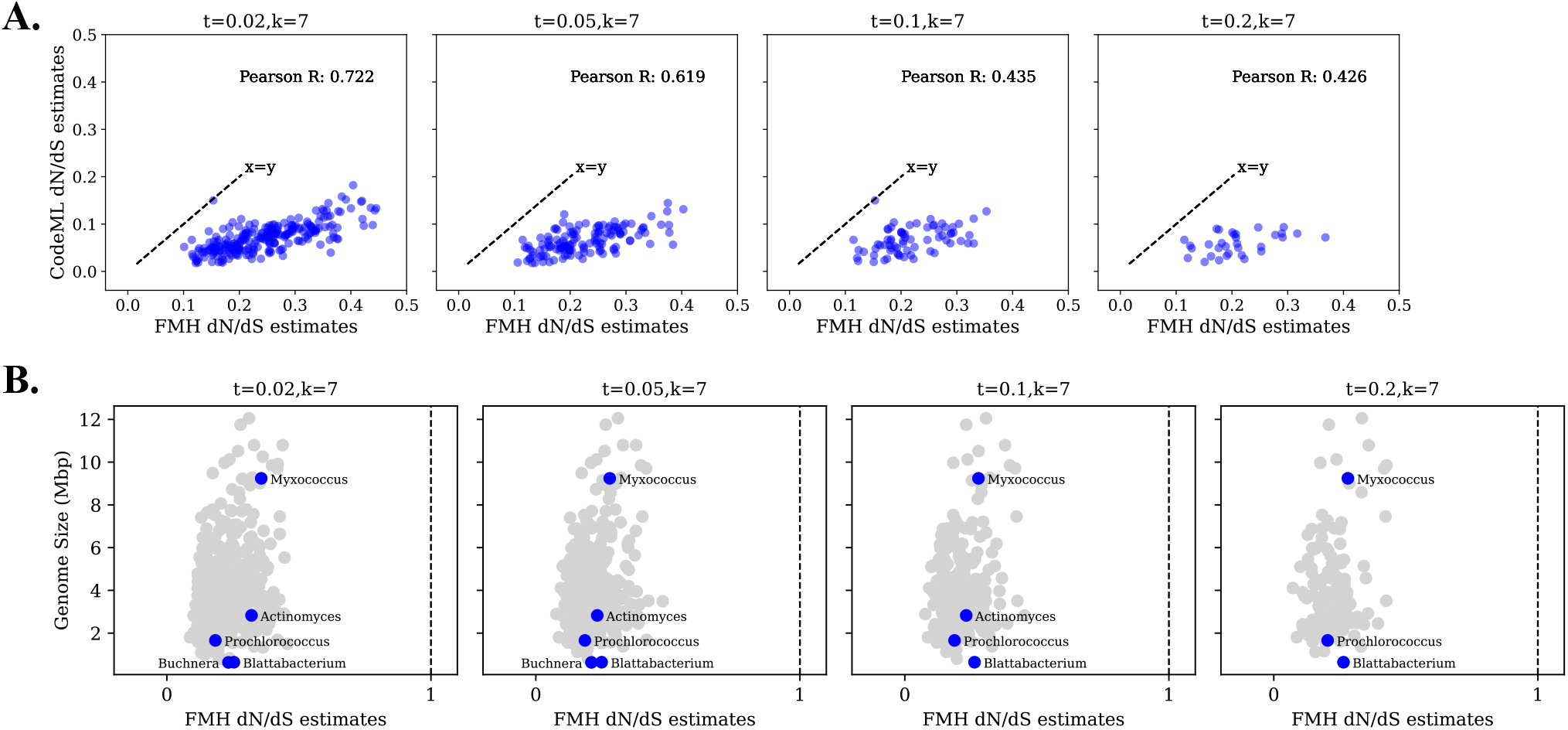
Applied *FracMinHash d*_*N*_/*d*_*S*_ on GTDB database. Estimations are based on a *k*_*aa*_=7 and *s*=500 across different containment thresholds of 0.02, 0.05, 0.1, and 0.2. Parameters Filtered values were based on a *pS* threshold of 0.05 and *pN* of 0.23. Data points are blue to indicate negative selection and neutral selection is indicated by a dotted vertical line at 1. A) Comparison of *FracMinHash d*_*N*_/*d*_*S*_ to an implemented maximum likelihood model, CodeML. Pearson R correlations are included. B) Evaluating interpretations made by *FracMinHash d*_*N*_/*d*_*S*_ when compared to genus genome size.

Although genomic *d*_*N*_/*d*_*S*_ overestimation is being observed by *FracMinHash d*_*N*_/*d*_*S*_, the inference of negative selection is correctly being made. To evaluate whether interpretation of results is affected, we attempt to reproduce the same interpretations made by from Martinez-Gutierrez (2022) focusing on the following genera: *Buchnera, Blattabacterium, Prochlorococcus, Actinomyces*, and *Myxococcus* (**Figure 4B**). As Martinez-Gutierrez (2022) observed, genera that have undergone extensive diversity loss—such as *Buchnera* and *Blattabacterium*—tend to exhibit higher *d*_*N*_/*d*_*S*_ values. In contrast, genera like *Prochlorococcus*, which inhabit stable open ocean environments maintaining genetic information, show lower *d*_*N*_/*d*_*S*_ values as expected. *Actinomyces* and *Myxococcus* have larger genomes and elevated *d*_*N*_/*d*_*S*_ values supporting the suggestion that genomic expansion is not favorable under either strong or weak purifying selection. In summary, the comparison of FracMinHash *d*_*N*_/*d*_*S*_ to real-world data demonstrate that despite the overestimations of genomic *d*_*N*_/*d*_*S*_, biologically meaningful and consistent interpretations of evolutionary pressures can be provided using FracMinHash *d*_*N*_/*d*_*S*_ .

### 3.3 Biological significances

We explore the application of *FracMinHash d*_*N*_/*d*_*S*_ to investigate conserved genomic regions between archaea and bacteria, and expand on the biological insights gained from *d*_*N*_/*d*_*S*_ analysis of orthologs across these domains. Specifically, we focus on a genomic region shared between *Methanobrevibacter sp*. (3.2 Mb), a gastrointestinal archaeon, and *Candidatus Saccharibacteria* (720.4 kb), a member of the TM7 bacterial phylum commonly found in the oral [30] and gut [31] microbiome. The data were sourced from BioProject PRJNA657473, which involved metagenomic sequencing of the goat gastrointestinal microbiome [32].

Archaeal and bacterial species have established a symbiotic relationship where bacteria donate electrons to archaea, while archaea offer bacteria a stable environment [33]. Evolutionarily, archaeal and bacterial genomes are traditionally viewed distinct, with Archaea typically exhibiting more compact genomes, shorter gene lengths, and reduced intergenic regions [34]. However, comparative genomics has increasingly revealed functional and structural parallels, particularly in conserved regions that may reflect shared ancestry or ancient horizontal gene transfer.

We used hierarchical edge bundling visualizations to highlight evolutionary relationships and help pin-point clusters of proteins under constraint or potential molecular adaptation from a genomic perspective [6]. This approach is particularly valuable for genomes that are incompletely annotated, as it enables the discovery of evolutionary patterns and conserved regions even in less-characterized taxa. We further evaluated *d*_*N*_/*d*_*S*_ at genetic level by using multiple traditional *d*_*N*_/*d*_*S*_ models to annotated orthologs within a conserved genomic island, we assess signatures of molecular selection that may suggest functional retention or adaptation.

Together, this integrated strategy enables broad-scale comparative analysis, reveals potential gene functions, and deepens our understanding of archaeal-bacterial genome evolution across distinct niches of the microbiome.

#### Hierarchical edge bundling facilitates identity signatures of selection among a group of genomes

To delve into the application of *FracMinHash d*_*N*_/*d*_*S*_ and expand on what biological significances can be obtained from FracMinHash *d*_*N*_/*d*_*S*_ results obtained from the previous genera discussed by producing and studying hierarchical edge bundling trees. Hierarchical edge bundling enables a clear visualization of large pairwise evolutionary relationships by bundling adjacent genomes together to reduce clutter. Using the *ggraph* R package, we produced a hierarchical edge bundling figure of the pairwise FracMinHash *d*_*N*_/*d*_*S*_ estimations generated from the GTDB database (**Figure 5A**). We sampled 100 random pairwise FracMinHash *d*_*N*_/*d*_*S*_ estimations across archaeal and bacterial genomes arranged within their genus groups bundling them into an evolutionary network. Clades under elevated adaptive pressures (FracMinHash *d*_*N*_/*d*_*S*_ > 1) are observed, as well as those under strong purifying pressures (FracMinHash *d*_*N*_/*d*_*S*_ *<* 1). Further, selection between broad taxonomic groups are also observed revealing novel signatures of selection, which would be hard to spot using a heatmap or tree.

**Figure 5:**
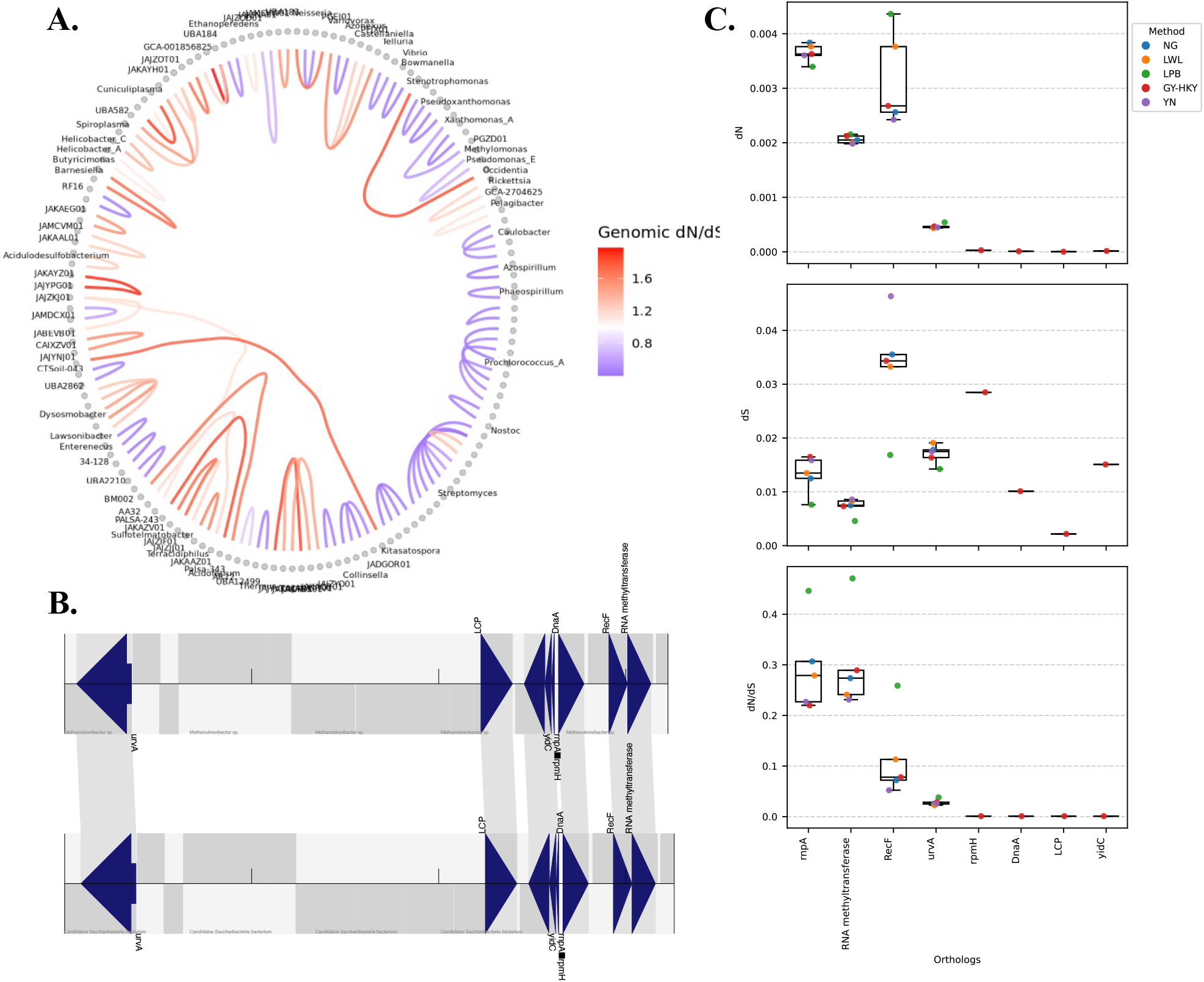
A. Hierarchical edge bundling visualization of pairwise selection between archaeal and bacterial genomes. B. Orthologous genes identified and annotated between *Methanobrevibacter sp*. and *Candidatus Saccharibacteria*. Genes including uvrA, LytR/CpsA/Psr (LCP), yidC, rnpA, rpmH, dnaA, and RNA methyltransferases appear in conserved order across both genomes. This arrangement suggests a vertically inherited genomic island maintained across archaeal and bacterial domains. Gene order and synteny support the functional conservation of this region. C. *d*_*N*_/*d*_*S*_ analyses indicate purifying selection across evaluated orthologs, with more synonymous substitutions than non-synonymous.

#### Genomic island between an archaeal and bacterial genome

Genes can be acquired by Archaea from Bacteria [35], raising the possibility that certain genomic regions—especially those resembling genomic islands—could be remnants of ancient or more recent horizontal gene transfer (HGT) events. HGT is an evolutionary process where genes are moved from one species to another [36]. Potential HGT events have been evaluated and identified using various methods including GC content [37, 38, 39], gene length, and gene synteny [40, 41, 42, 43, 44, 45]. Here, we investigate this hypothesis by focusing on a conserved genomic island-like region shared between archaeal and bacterial genomes.

We begin this evaluation by using *nucmer* for global alignment, the genomic sequences of the archaea and bacterial pairs of interest are aligned to identify regions potentially under selection (**Table 1**). Interestingly, we discovered genomic regions shared between archaeal and bacterial genomes. These shared regions were processed using in-house Python scripts to identify potential orthologs. Subsequently, we used *mummerplot* to visualize the alignments and highlight candidate regions for further investigation. One region of particular interest spanned 35,796 bp and was shared between two scaffolds (**Figure 5B**). These scaffolds were extracted and annotated using *prodigal* to identify coding regions and their corresponding protein sequences. Note, we used *prodigal* because protein sequences are required for *proteinortho* [46] when using the BLASTP option. Potential orthologs were identified using *proteinortho*, and because they were unannotated, we used *diamond* in combination with the Swiss-Prot database to assign UniProt IDs and infer protein function.

**Table 1:**
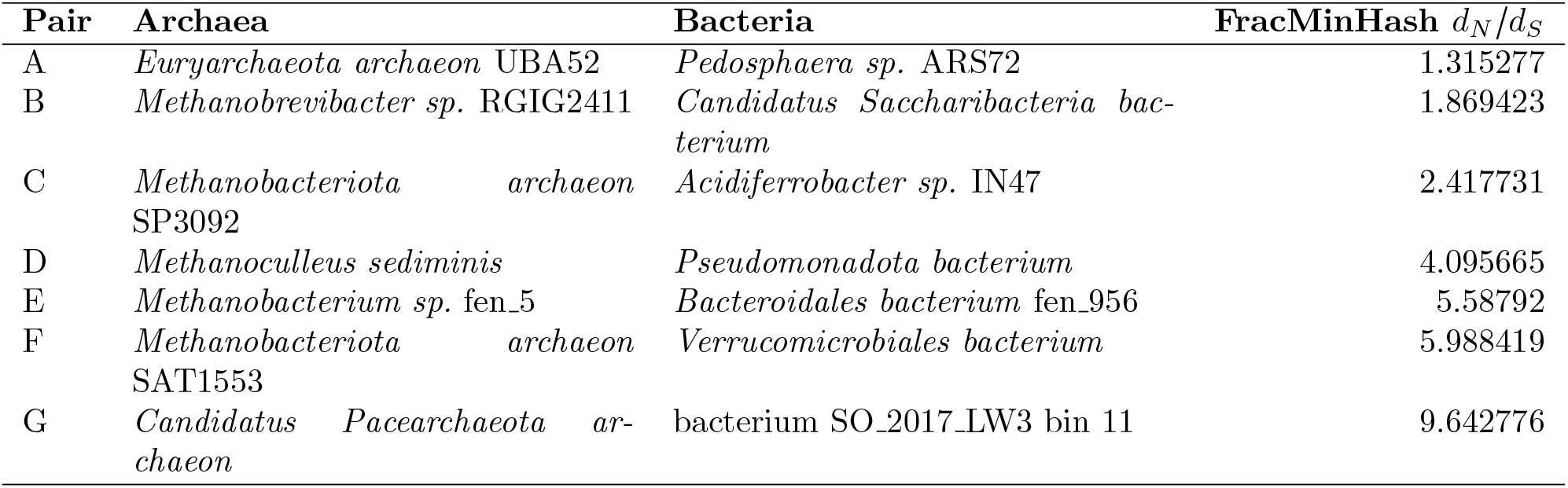
Genome pairs sorted by FracMinHash *d*_*N*_/*d*_*S*_ (least to largest). In this scenario the archaea is the reference. Here, we look at a FracMinHash *d*_*N*_/*d*_*S*_ between 1.1 and 10.

The proteins encoded within this region were located in close proximity and showed conserved gene order, therefore supporting the interpretation of this region as a genomic island. Conserved synteny across divergent species, particularly between *Methanobrevibacter sp*. and *Candidatus Saccharibacteria*, can reveal evolutionary relationships and potential HGT events. Synteny-based methods have been used to detect antimicrobial resistance [40, 41, 43], virulence factors [42, 43], and record strain tracking [44, 45]. A noteworthy finding in this study is the conserved genomic arrangement of the identified genes (**Figure 5B, Supplemental Table ??**). This conservation of gene order, especially across distantly related lineages, may indicate a horizontal gene transfer event that preserved not only gene content but also local synteny.

An additional sign of HGT is GC content, a measure of DNA stability where more GC in a sequence indicates stability [37, 38, 39]. According to publicly available annotations, *Methanobrevibacter sp*. contains 3,233 protein-coding genes with a GC content of 32.5%, while Candidatus Saccharibacteria has 693 protein-coding genes with a GC content of 41.0%. When evaluating the GC content of the orthologous genes of interest, we found a range of 28.6%–40.3% (mean = 34.9%) and 28.8%–40.6% (mean = 34.8%) for each genome, respectively. The distribution of GC content (**Supplemental Figure ??**) shows a moderate shift to the left compared to the overall distribution in *Candidatus Saccharibacteria bacterium* RGIG2249, thus supporting the possibility of foreign gene origin.

We also assessed the distributions of gene length, another clue for potential HGT. We compared the distribution of gene lengths in whole genomes versus the orthologs of interest (**Supplemental Figure ??**). In this assessment, no significant differences were found in the distributions, suggesting this genomic-island may have adapted to the protein-coding lengths of their host genomes, but further study is needed.

Additionally, we evaluated the distribution of this genomic region in other samples using NCBI and LoganSearch. By using NCBI, we searched a random *Methanobrevibacter sp*. RGIG5977 strain and by using *ProteinOrtho*, two hits for the UvrA protein were reported—each 2889 bp long with 37% GC content—located on separate contigs (**Table 2**), suggesting a recent intragenomic duplication event. This process was repeated on a *Methanobrevibacter sp*. MAG001 strain drawn from a different BioProject (PRJNA647157, a sample obtained from a Tibetan pig gut), where a single ortholog was found, UvrA (Accession WP 178787431.1, 2,886 bp, GC content of 34%, found in Contig NZ JAEDCH010000024.1). Interestingly, UvrA and UvrB have been shown to suppress spurious recombination [47]. No other genes from the genomic island were reported as orthologous to *Methanobrevibacter sp*. MAG001.

**Table 2:** UvrA duplication in *Methanobrevibacter sp*. RGIG5977.

| Accession | Location |
| --- | --- |
| NZ_JAFWQL010000013.1 | 11986–14874 |
| NZ_JAFWQL010000007.1 | complement(22592–25480) |

Repeating this for a *Candidatus Saccharibacteria bacterium* strain, we find all of the orthologs of this genomic region in *Candidatus Saccharibacteria bacterium* oral taxon 955 from a different BioProject (PRJNA981146) (**Table 3**); however, synteny is not as strictly maintained as in *Methanobrevibacter sp*. RGIG2411 and *Candidatus Saccharibacteria bacterium* RGIG2249. For example, RNA methytransferase is on the complement strand in *Candidatus Saccharibacteria bacterium* oral taxon 955. Additionally, the GC content is elevated compared to *Methanobrevibacter sp*. RGIG2411 and *Candidatus Saccharibacteria bacterium* RGIG2249 with an average GC content of 0.47%. However, *Candidatus Saccharibacteria bacterium* oral taxon 955 containing these genes while other strains do not suggests evidence that potentially the genomic island found in *Methanobrevibacter sp*. RGIG2411 was acquired from *Candidatus Saccharibacteria bacterium* RGIG2249.

**Table 3:** Genes of genomic coordinates found in *Candidatus Saccharibacteria bacterium* oral taxon 955. Information includes the length of the protein-coding sequence, and GC content

| Protein family | Coordinates | Length (bp) | GC content |
| --- | --- | --- | --- |
| UvrA | complement(628690--631503) | 2814 | 0.4737 |
| LCP | 174844–176511 | 1668 | 0.4718 |
| YidC | complement(860649--861578) | 930 | 0.4140 |
| RnpA | complement(861639--861983) | 345 | 0.4319 |
| rpmH | complement(862040--862177) | 138 | 0.5580 |
| DnaA | 1–1350 | 1350 | 0.4696 |
| RecF | 2720–3751 | 1032 | 0.4293 |
| RNA methyltransferase | complement(363560--364864) | 1305 | 0.4881 |

We further used LoganSearch [48], a *k*-mer-based search engine that facilitates querying sequences against metagenomic samples, to find samples that have this genomic region at or above 80% nucleotide identity. The region appeared in multiple gastrointestinal sites (Ileum, Jejunum, Rectum, Cecum, Duodenum, Rumen, Reticulum, and Colon) within the same Bioproject, which shows that it’s unlikely to be a bioinformatics artifact and pertains to the goat-host as it showed up in multiple locations.

In summary, our findings provide strong evidence that this genomic island was acquired via a recent HGT event between *Methanobrevibacter sp*. RGIG2411 and *Candidatus Saccharibacteria bacterium* RGIG2249. Although average gene length shows little divergence and GC content varies across both genomes and their orthologous groups, we observed conserved synteny and clear orthologous relationships. Moreover, the functional coherence and structural conservation of the gene cluster, which was not found in other strains, further support this recent HGT event.

#### Synonymous substitutions occur at higher rates than non-synonymous substitutions

To investigate selective pressure on the conserved orthologs within the shared genomic region, we evaluated multiple traditional *d*_*N*_/*d*_*S*_ models using the program *KaKs_Calculator* [49] (**Figure 5C**). Orthologous coding sequences between *Methanobrevibacter sp*. RGIG2411 and *Candidatus Saccharibacteria bacterium* RGIG2249 were aligned using *ClustalW* [50] and converted to the required *AXT* file format. We tested five traditional *d*_*N*_/*d*_*S*_ models on these alignments: NG86, LWL, LPB, GY-HKY, and YN.

The estimated *d*_*N*_/*d*_*S*_ values were below 1, indicating more synonymous mutations and suggesting constraining selection. Among these, four proteins stood out with slightly elevated *d*_*N*_/*d*_*S*_ values (though still <1): Rnp4, Class I-like SAM-binding methyltransferase, RNA m5U methyltransferase, RecF, and UvrA (a member of the ABE transporter family). Although model estimates varied slightly, values were consistently much lower than values, reinforcing the conclusion that these genes are under strong purifying selection and are likely functionally important and conserved. A small subset of proteins, including ribosomal protein bL34, DnaA, LytR/CpsA/Psr (LCP) family, and OXA1/ALB3/YidC Type 1 family, produced *d*_*N*_/*d*_*S*_ values only under the GY-HKY model; other models failed to produce results for such closely related sequences (**Supplemental Table ??**). In these cases, values were elevated while values approached zero, further supporting strong functional conservation.

Among the protein-coding genes identified, a potential RNA methyltransferase sharing a domain related to the Class I-like SAM-binding methyltransferase and RNA m5U methyltransferase families were found. These families are broadly conserved and often uncharacterized, as evidenced by the numerous uniprot hits reported by *diamond*. Methyltransferases play important roles in microbial metabolism within the gastrointestinal tract. Interestingly, *Methanobrevibacter sp*. RGIG2411 have been found to be more abundant in the small intestine, suggesting a specialized role in anaerobic digestion [32]. In anaerobic digestion, organic waste is converted to gases, such as methane, and have been an area of research due to their gas products potential in renewable energy [51]. Specifically, methanogenesis, the production of methane, is part of anaerobic digestion and use the methyltransferases to transfer methyl groups and are found involved in methanogenesis [52]. In contrast, the functional roles of methyltransferases in *Candidatus Saccharibacteria bacterium* RGIG2249 remain unclear. However, previous work has proposed horizontal gene transfer of methyltransferase genes, particularly RNA m5U methyltransferases, from bacteria to archaea [35]. Therefore, further characterization of these methyltransferase-encoding regions in *Methanobrevibacter sp*. RGIG2411 and *Candidatus Saccharibacteria bacterium* RGIG2249 is required to elucidate the functional significance and evolutionary history of these protein-coding regions. These regions correspond to NCBI IDs IJH63 12540 (*Methanobrevibacter sp*. RGIG2411) and IJG01_03160 (*Candidatus Saccharibacteria bacterium* RGIG2249).

Additional orthologs of interest were also evaluated. A coding sequence resembling the LytR/CpsA/Psr (LCP) family was identified, showing moderate similarity to BrpA (UniProt: BRPA_STRMU) from *Streptococcus mutans*, a dental pathogen [53]. BrpA has been implicated in regulating biofilm formation promoting *S. mutans* virulence [54, 55]. The BrpA-like sequences identified correspond to NCBI IDs IJH63 12500 (*Methanobrevibacter sp*. RGIG2411) and IJG01_03120 (*Candidatus Saccharibacteria bacterium* RGIG2249). Given the oral cavity is the first site of microbial interaction during digestion, the connection between oral and gut microbial genes warrants further investigation. Similarly, another coding sequence homologous (IJH63_12510 and IJG01_03130 for *Methanobrevibacter sp*. RGIG2411 and *Candidatus Saccharibacteria bacterium* RGIG2249, respectively) to YidC, part of the OXA1ALB3YidC family, Type 1 subfamily involved in protein insertions and can serve as an antibiotic target [56, 57]. The coding regions within this genomic island contain domains associated with metabolic functions, suggesting it may constitute a metabolic island [37]. However, the presence or role of protein families and their direct evidence of expression or function in the context of the gut can be further explored. For other orthologs, no clear links to gut-related functions were found. These cases highlight the limitations of current annotations and the need for further functional studies to determine their biological significance.

## 4 Discussion

In this study, we present and evaluate *FracMinHash d*_*N*_/*d*_*S*_, an alignment-free approach for estimating genomic *d*_*N*_/*d*_*S*_ ratios (FracMinHash *d*_*N*_/*d*_*S*_) with potential applications to large-scale genomic and metagenomic data. Simulation results demonstrate that *FracMinHash d*_*N*_/*d*_*S*_ achieves strong agreement with traditional *d*_*N*_/*d*_*S*_ models (e.g., NG86), particularly for longer sequences and moderate *k*-sizes (*k*_*aa*_ = 5–7), with correlations exceeding a Pearson *R* of 0.90. While larger *k*-sizes reduce correlation, they improve runtime and disk efficiency, highlighting the method’s scalability. Even with scale factors as high as 1000, meaningful estimates can still be obtained.

When applied to real-world genomic data, including mutated biological sequences and previously analyzed genera, *FracMinHash d*_*N*_/*d*_*S*_ successfully recapitulates biologically meaningful patterns of selection. Notably, it correctly infers negative selection despite moderate overestimation of *d*_*N*_/*d*_*S*_ values. In taxa such as *Buchnera* and *Blattabacterium*, where genome reduction is prominent, elevated *d*_*N*_/*d*_*S*_ values are consistent with previous findings. Conversely, low *d*_*N*_/*d*_*S*_ values in *Prochlorococcus* align with expectations for taxa in stable environments, while *Actinomyces* and *Myxococcus* exhibit patterns reflecting the impact of genomic complexity on selection.

We further applied traditional *d*_*N*_/*d*_*S*_ models to a conserved genomic island shared between *Methanobrevibacter sp*. RGIG2411 and *Candidatus Saccharibacteria bacterium* RGIG2249. Our findings reveal signatures of purifying selection and conserved gene order, suggesting an HGT event of functionally important genes. These results support the view that, despite the evolutionary divergence between archaea and bacteria, regions of conserved function and ancestry can be detected and meaningfully interpreted using *FracMinHash d*_*N*_/*d*_*S*_.

Although overestimations were observed with *FracMinHash d*_*N*_/*d*_*S*_, the resulting *d*_*N*_/*d*_*S*_ interpretations remained biologically consistent when compared to both a counting-based model (i.e., NG86) and a maximum-likelihood model (i.e., codeML). The performance of our model was not evaluated in the presence of recombination, which is known to affect *d*_*N*_/*d*_*S*_ estimates. Similar to NG86, *FracMinHash d*_*N*_/*d*_*S*_ is based on a simplified model and was not designed to account for recombination or to distinguish between transitions and transversions, as more complex *d*_*N*_/*d*_*S*_ models do [5].

Moreover, to assess the applicability of *FracMinHash d*_*N*_/*d*_*S*_ on raw metagenomic reads, we evaluated its performance using six-frame translations of simulated data to estimate FracMinHash *d*_*N*_/*d*_*S*_ (**Supplemental Figure ??**). This approach enables the translation and sketching of unannotated genomic sequences, such as complete genomes or raw sequencing reads. However, *d*_*N*_/*d*_*S*_ estimates obtained through six-frame translation showed poor performance on the simulated datasets. In contrast, estimates derived from annotated coding sequences generated with *prodigal* demonstrated significantly improved accuracy, indicating that applying gene prediction prior to running *FracMinHash d*_*N*_/*d*_*S*_ enhances *d*_*N*_/*d*_*S*_ estimation. Therefore, developing a theoretical framework to accurately estimate dN/dS directly from six-frame translations would be a valuable advancement for analyzing raw metagenomic data.

Overall, this work highlights the potential of FracMinHash *d*_*N*_/*d*_*S*_ as a scalable, computationally efficient tool for evolutionary analysis across diverse taxa, even in the absence of high-quality alignments or complete annotations. It enables comparative genomics, evolutionary inference, and functional interpretation across both synthetic and complex biological datasets, and may potentially help identify genomic signatures useful for strain tracking, diagnosis, treatment, and drug development.

## Acknowledgements

We thank Jinglin Feng on the contributions for the Hierarchical Edge Bundling. Research reported in this publication was supported by NIGMS of the National Institutes of Health under award number R01GM146462. The content is solely the responsibility of the authors and does not necessarily represent the official views of the National Institutes of Health.

